# Causal approach to environmental risks of seabed mining

**DOI:** 10.1101/2021.02.21.432138

**Authors:** Laura Kaikkonen, Inari Helle, Kirsi Kostamo, Sakari Kuikka, Anna Törnroos, Henrik Nygård, Riikka Venesjärvi, Laura Uusitalo

## Abstract

Seabed mining is approaching the commercial mining phase across the world’s oceans. This rapid industrialization of seabed resource use is introducing new pressures to marine environments. The environmental impacts of such pressures should be carefully evaluated prior to permitting new activities, yet observational data is mostly missing. Here, we examine the environmental risks of seabed mining using a causal, probabilistic network approach. Drawing on a series of interviews with a multidisciplinary group of experts, we outline the cause-effect pathways related to seabed mining activities to inform quantitative risk assessments. The approach consists of (1) iterative model building with experts to identify the causal connections between seabed mining activities and the affected ecosystem components, and (2) quantitative probabilistic modelling to provide estimates of mortality of benthic fauna in the Baltic Sea. The model is used to evaluate alternative mining scenarios, offering a quantitative means to highlight the uncertainties around the impacts of mining. We further outline requirements for operationalizing quantitative risk assessments, highlighting the importance of a cross-disciplinary approach to risk identification. The model can be used to support permitting processes by providing a more comprehensive description of the potential environmental impacts of seabed resource use, allowing iterative updating of the model as new information becomes available.

## 1. INTRODUCTION

The oceans are facing increasing pressures from human activities. The intensified use of marine space and resources is embodied both through expansion of existing activities (Halpern et al. 2015), and creating new industries for marine resource use (Voyer et al. 2018; Winther et al. 2020). To ensure sustainable development of maritime activities, the impacts of new types of activities should be carefully evaluated prior to permitting them (Borja et al. 2016). Seabed mining is one of the rapidly emerging sectors promoted to support resource sufficiency, with especially the deep seabed presented as a new frontier for resource extraction (Hein et al. 2013). However, dealing with impacts of activities that do not take place yet means that there is no observational data on the impacts, with high uncertainties on both the implementation of the activity and its consequences for the environment. This uncertainty creates a challenge to estimate the impacts in a way that is scientifically robust, while accounting for the knowledge gaps and scarcity of data to support decision-making.

Current plans for mining are outlined both in shallow continental shelf areas and the deep sea, encompassing areas within national jurisdiction of sovereign states and the international seabed in the ‘Area’ (Miller et al. 2018). While most initiatives are still at an exploratory stage, the increasing need for raw materials is pushing countries to consider where to get their mineral resources in the future (Vidal et al. 2017).

Seabed mining will likely affect all levels of marine ecosystems, including the water column and the seafloor (Boschen et al. 2013; Kaikkonen et al. 2018; Miller et al. 2018). The potential environmental impacts of mining have been addressed in an increasing number of studies (Miljutin et al. 2011; Jones et al. 2017; Orcutt et al. 2018; Simon-Lledó et al. 2019). Even with valuable data from these experiments, the impact studies conducted to date offer a scattered view of the environmental impacts, with no attempts to synthesize impacts to support an operational assessment. It is further uncertain to what extent the empirical disturbance studies succeed in scaling up to industrial mining operations(Jones et al. 2017).

Environmental risk assessment (ERA) is a process aiming to evaluate the different possible outcomes following human activities (Burgman 2005). A risk in this context is defined as any unwanted event (here ‘impact’) and its probability. Currently, most ERAs build on estimating ecosystem responses to pressures based on vulnerability of the environment through semi-quantitative scoring instead of the activity itself (Stelzenmueller et al. 2015; Washburn et al. 2019; Quemmerais-Amice et al. 2020), and as such are not well suited for describing different possible combinations of outcomes from new untested activities. By assuming additive relationships of pressures, these approaches often neglect the synergistic and antagonistic effects of pressures (Halpern and Fujita 2013).

A broader appreciation of the risks in the context of new maritime activities thus calls for improved systems thinking, structured approaches, and integration of knowledge from multiple sources and disciplines (Holsman et al. 2017). Updating of prior knowledge is important to evaluate to what extent new studies could decrease the uncertainties. A first step towards a comprehensive view of the risks stemming from seabed mining activities requires identifying the sources of changes in the environment, affected ecosystems components, and any further variables associated with these.

Drawing on the recognition of causes and effects, causal chains or networks offer a systematic method to study environmental impacts (Perdicoúlis and Glasson 2006). By describing the factors affecting the state of the system in as much detail as possible, causal networks enable evaluating multiple scenarios and improve understanding of the underlying mechanisms in the studied system (Pearl 2009). When applied in environmental management, causal approaches have been shown to be useful in policy interventions and management (Carriger et al. 2016; Carriger et al. 2018).

Bayesian networks (BNs) are graphical models that represent a joint probability distribution over a set of variables and provide an alternative to commonly used scoring procedures in ERAs (Pearl 1986; Kaikkonen et al. 2021). In BNs, the strength of each connection between variables is described through conditional probabilities. As probabilistic models, the result of a BN is not a single point estimate, but a probability distribution over the possible values of each variable, allowing estimating not only the most likely outcome, but also the uncertainty associated with the estimates (Varis et al. 1990; Fenton and Neil 2012). BNs can thus be used to synthesize outcomes of multiple scenarios by evaluating possible combinations of events and weighting them according to how likely they are. Given their modular structure, they can be used to support integrative modelling and can accommodate inputs from multiple sources, including simulations, empirical data, and expert knowledge (Uusitalo 2007; Helle et al. 2020).

Here, we describe an approach for integrating expert knowledge into a causal risk assessment for seabed mining. We use the Baltic Sea as an example to test our approach, as mining iron-manganese nodules has already been tested in an industrial setting in this area (Zhamoida et al. 2017) and the ecosystem components and food web structure are well studied (Yletyinen et al. 2016; Reusch et al. 2018; Törnroos et al. 2019). Given the number of ongoing seabed mining initiatives and attempts to quantify impacts, the aim of this work is to provide a framework that allows combining information from multiple sources by bringing ecological information to risk analysis while explicitly addressing uncertainty. To move towards a quantitative risk assessment, we demonstrate the use of BNs in an operational setting and discuss needs for a quantitative ERA in the context of emerging maritime activities.

## 2. CASE STUDY BACKGROUND

Our case study deals with ferromanganese (FeMn) concretion removal in the northern Baltic Sea. The Baltic Sea is characterized by low species richness compared to many marine areas, and the food web structure and ecological traits characterizing major taxa have been well described (Törnroos and Bonsdorff 2012). Due to the relatively shallow depth of the Baltic Sea, the extraction activity is to some extent comparable to sand and gravel extraction and would likely be performed by suction hopper dredging (Zhamoida et al. 2017).

In our study scenario, mineral extraction is restricted to areas with a minimum depth of 40 meters, assuming regulatory limits of such activities below the aphotic zone (Kostamo 2021). The densest occurrences of FeMn concretions in Baltic Sea are also found below these depths (Kaikkonen et al. 2019). We assume that extraction is performed in a zig-zag pattern in a limited extraction area of 1 km^2^ and it removes all concretions in the path of the suction head (Fig. 1). Here we assume homogeneous impacts on the areas that are not subject to direct extraction, although in reality the spatial footprint of impacts is dependent on the particle movement and distance of a point from the extraction area (Smith and Friedrichs 2011; Spearman 2015). Risks related to operating the vessels and impacts during transportation are not considered, as they are well addressed in other studies (Kulkarni et al. 2020).

**Figure 1.**
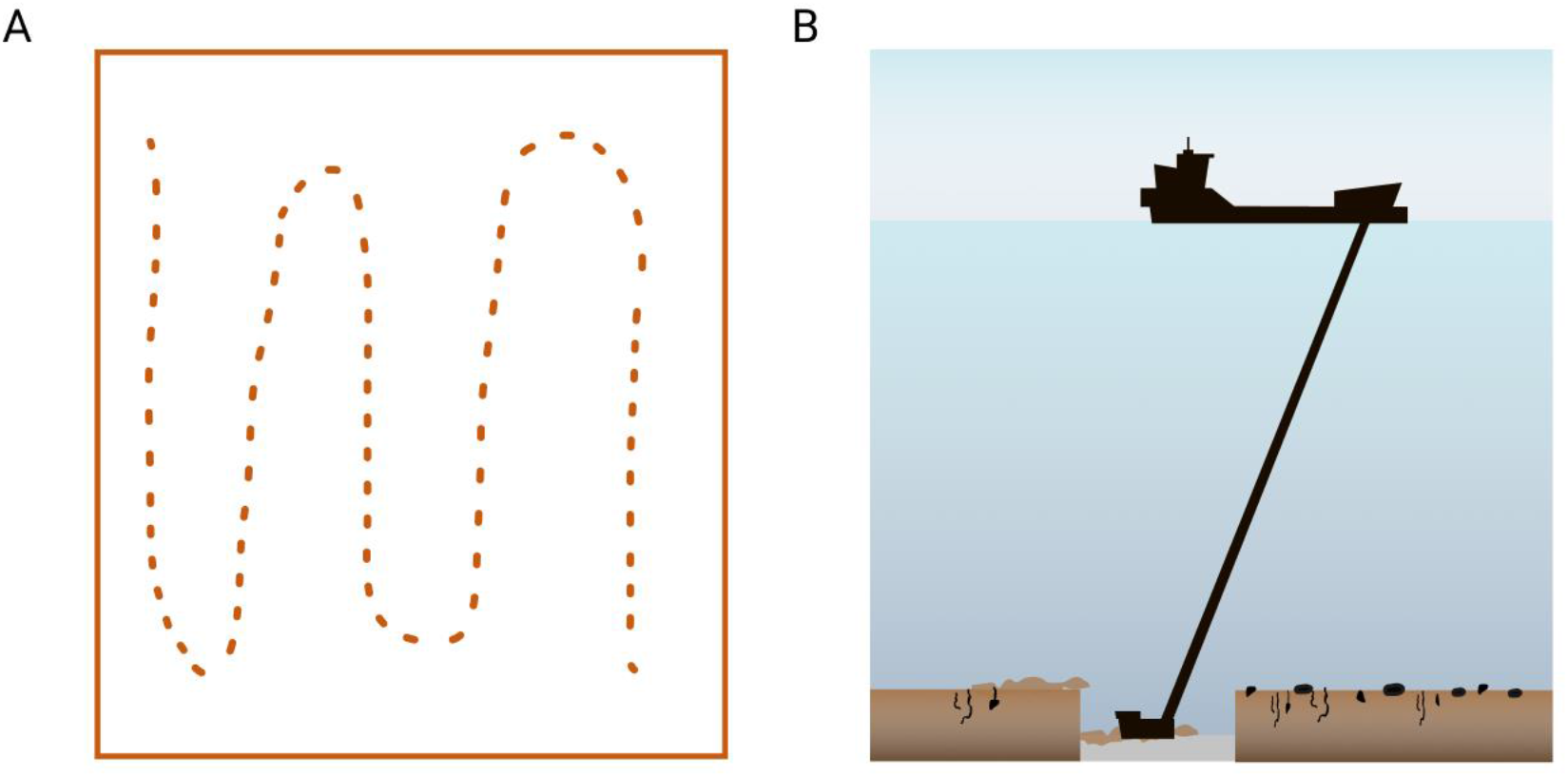
A) Plan view and B) profile view of mining a 1 km^2^ mining block. The dotted lines in panel A illustrate the extraction pattern of the mining device in a discrete block with FeMn concretions.

## 3. METHODS

We apply a 3-step approach for working together with experts to create a model that summarizes the causal connections in the system and enables providing quantitative risk and uncertainty estimates (Fig. 2). The first step consists of mapping the relationships between key drivers and ecosystem responses with experts in semi-structured interviews. The use of structured methods for expert elicitation has been highlighted in recent years, and here we follow a modified version of the IDEA (Investigate-Discuss-Estimate-Aggregate) protocol that consists of both individual and aggregated assessments from experts (Burgman 2016; Hemming et al. 2018). Although the method is designed for quantitative estimates, here we use it only for qualitative causal mapping to test a structured approach for more comprehensive interviews. In the second step, a combined model structure is created and reviewed by the experts in an iterative manner until a satisfactory model structure was obtained. The final step consists of quantifying the magnitude of the ecosystem impacts through conditional probabilities.

**Figure 2.**
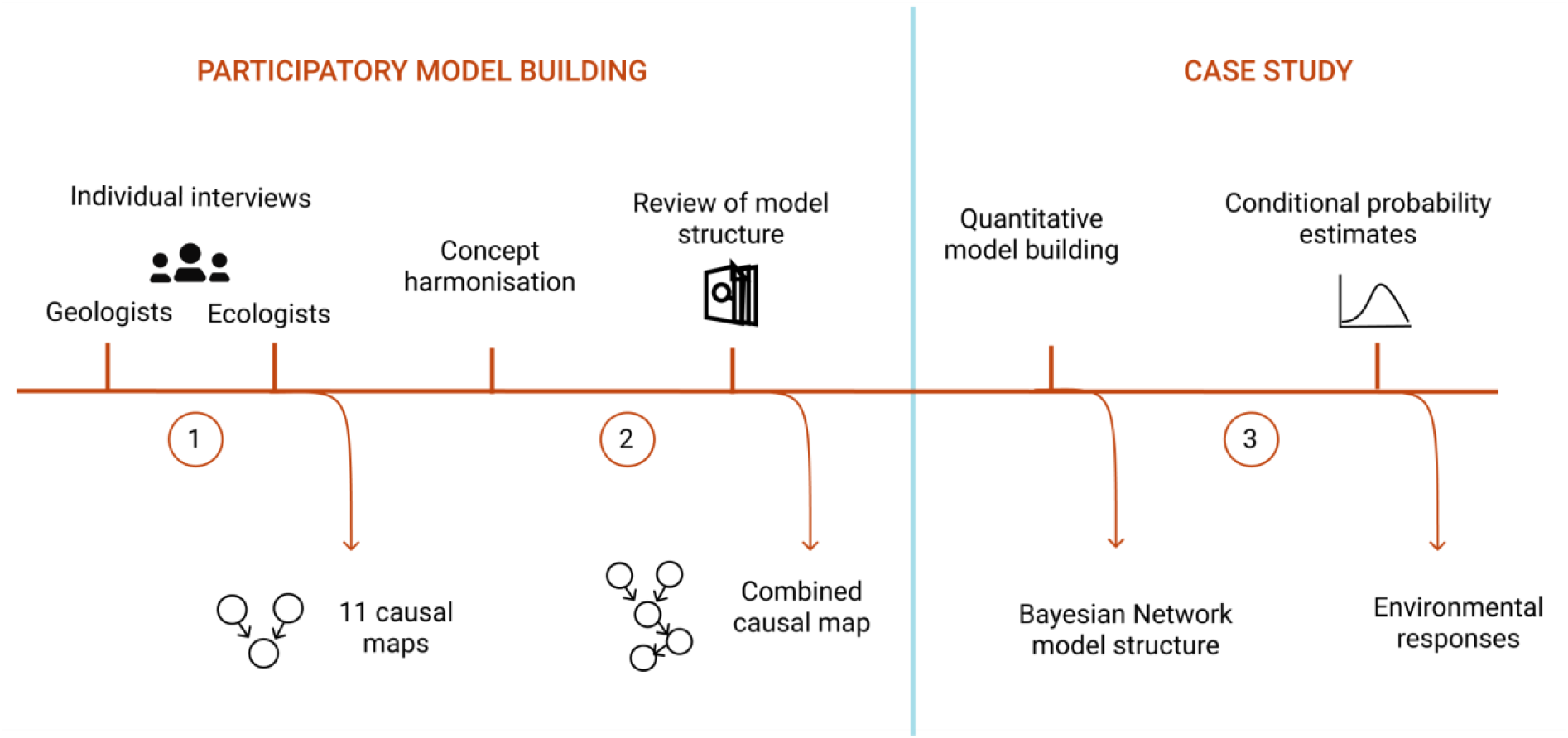
Conceptual figure of the modelling process summarizing the activities within the proposed approach (upper panel) and four main outcomes (lower panel).

### 3.1 Step 1: Expert interviews

Framing the system and the connections between variables was performed as a causal mapping exercise with a multidisciplinary group of experts. The aim of causal mapping is to explore an individual’s view on a system under different scenarios by detailing the causes and effects. In an ERA context, this step constitutes the risk identification stage (Suter II 2016). Experts were recruited through snowball sampling by consulting researchers in different fields of marine sciences. To attain a diverse sample and sources of knowledge, we sent invitations to experts representing varying backgrounds in different institutes. The final list of experts participating in the study included 11 experts from universities in Finland and Sweden, governmental research institutes, as well as intergovernmental organizations working on the Baltic Sea (ICES, HELCOM).

The causal mapping exercise was conducted through semi-structured interviews. We used individual interviews, as group interviews can be dominated by a small number of individuals (Martin et al. 2012), and experts’ judgments can be influenced by their peers (O’Hagan et al. 2006). Gradual elicitation allowed us to evaluate when a sufficient number of experts had been interviewed by monitoring when the number of variables no longer increased with the addition of new experts.

Semi-structured interviews were held at a location chosen by the interviewee or via an online connection. For face-to-face interviews, causal maps were drawn on paper, whereas in online interviews maps were constructed using an online drawing tool. All interviews were recorded with consent from the interviewee.

At the beginning of each interview, participants were introduced to the use of causal networks. Each expert was presented with the same scenario of the mining activity and the changes in the environment arising from the activity, noted as pressures (Table 1). Details on how mining would likely be carried out were drawn from literature and informal consultation with experts in geology and mineral resource extraction.

**Table 1.** Physicochemical changes in the environment (pressures) arising from mining used as a starting point in causal mapping with experts.

| Pressure type | Description and references |
| --- | --- |
| Nodule removal | FeMn concretion removal from a mining block. Contributes to loss of hard substrate on otherwise soft seabed. |
| Modification of seafloor substrate type | Measure of changes in the sediment environment, including changes in:<br>· Grain size<br>· Sediment porosity<br>· Sediment compaction |
|  | · Organic enrichment<br>· Pore water composition<br>· Oxygen penetration depth |
| Modification of seafloor topography | Changes in seafloor topography following extraction activities (Zhamoida et al. 2017). |
| Sediment dispersal in the water column | Total suspended solids concentration near the surface or in the water column both from the processing return and mining tool operation (Spearman 2015). |
| Sediment dispersal near seafloor | Total suspended solids concentration near the seafloor resulting from the processing return and mining tool operation (Sharma et al. 2001). |
| Release of nutrients from the sediment | Release of soluble nutrients from the sediment plume to the seabed water column (Jones and Lee 1981; Lohrer and Wetz 2003). |
| Release of toxic substances from the sediment | Release of contaminants from the sediment plume to the water column (Simpson and Spadaro 2016; Hauton et al. 2017; Couvidat et al. 2018). |
| Underwater noise | Noise from the mining operation, including extraction of the substrate and vessel operations (Robinson et al. 2011; Theobald et al. 2011). |

The first three interviews were held with marine geologists with experience in underwater mining technology. These interviews were used to adjust the pressures identified in a literature review and to identify environmental parameters and operational factors likely to affect the magnitude of the physiochemical changes arising from mining (Table 1). These variables form the core of the model by describing the basic processes related to mining.

To explore the ecological impacts arising from these pressures, the following eight interviews were conducted with marine ecologists. Each expert was presented with the same scenario of the mining activity and the physicochemical pressures identified in the first phase with the geologists (Table 1). The experts were then asked which ecosystem components they think will be affected by these pressures. Whenever possible, experts were asked to rate the strength of the causal connection on a 1–3 scale. As the number of individual species even in the relatively species-poor Baltic Sea is too high to include in one model, we reduced this complexity by asking experts to address the affected organisms through the functional traits that would differentiate the effects on these organisms.

Experts were given unlimited time to complete the causal map and were informed that they may modify the causal map after the interview. After each interview (approximately 2–3 hours each), the causal maps were digitized, and the resulting maps were sent to the experts for verification.

### 3.2 Step 2: Combining causal maps

To obtain a comprehensive view of the environmental impacts of mining, the individual causal maps were combined into one causal network. To do this, we coded the connections between variables in the individual causal maps to adjacency matrices using the assigned link strengths whenever available. Prior to combining the maps, variables were harmonized and combined so that similar concepts were grouped under one variable. For instance, the terms “polychaetes”, “annelids”, and “worms” were grouped under ‘mobile infauna’ (see Table S1 in Supporting Information for full details of individual maps).

The final list of functional groups was compiled from the traits and taxa mentioned in the expert interviews and groupings used in other studies(Hewitt et al. 2018) based on the expected response of organisms to the pressures caused by mining so that the traits characterize differential responses in the organisms. Here, traits are treated as binary variables, although most species express a variety of traits (Villnäs et al. 2018).

While elicitation of individual causal maps has been explored in depth in literature (Özesmi and Özesmi 2004; LaMere et al. 2020), there is little guidance on how to systematically combine diverse variables into one consensus map. In this work, all non-redundant variables and connections were included in the combined network. To ensure that the combined map represented the views of the experts involved in the model framing, experts had the possibility to comment on the network structure in an open online document presented both in the form of a graph and a table. At this stage, the document and the comments were visible to all experts.

### 3.3 Step 3: Bayesian Network model development

The final causal network was used to develop a probabilistic Bayesian network (BN) to provide quantitative estimates of the ecological consequences of mining to ecosystem components under different mining scenarios. In this work, we quantified only a sub-model of the complete causal network focusing on three groups of benthic fauna: sessile filter feeding epifauna, mobile epifauna, and burrowing infauna. The BN model was developed from variables describing these three benthic faunal groups, the main pressures affecting them, and any intermediate variables between them in the combined causal network. To reduce complexity of the model in terms of spatial and temporal dimensions of the impacts, we restricted the model to account only for the acute impacts within a spatially discrete mining block as defined in the case study description (Fig. 1). Discrete variable states were drawn from literature and expert views. We use relative descriptions of pressures with relation to ambient conditions (e.g. low-high). To evaluate the model structure, we conducted a point-by-point walkthrough of the model with external experts in marine ecology and geology who had not participated in the model building.

To quantify the magnitude of impacts between the pressures and the benthic faunal groups, we modelled the BN as an expert system, meaning that no empirical data is directly incorporated in the model. We used the graphical interface provided open source Application for Conditional probability Elicitation (ACE) (Hassall et al. 2019) to initialize the conditional probability tables (CPTs) with one expert in geology and one benthic ecologist. The application provides a starting point for defining the overall shape of a conditional probability distribution, which is done by ranking the direction and magnitude of the parent nodes on the child node and populating the table through a scoring algorithm (Hassall et al. 2019). The scoring system considers that all variable states can be placed on an equally spaced linear scale.

To assess probabilities of the impacts of direct pressures on benthic fauna, the CPTs initialized with the ACE application were evaluated and adjusted in a second session with another benthic ecologist. The total mortality of benthic fauna within a discrete block and one moment in time comprises the direct mortality from extraction of sediment and mineral concretions, and the indirect mortality of the remaining fauna that are exposed to the pressures from the extraction activity. The probability of total mortality of benthic fauna was thus calculated as:

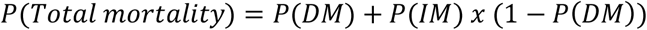

where the term IM x(1-DM) accounts of the proportion of fauna remaining after direct extraction. We applied numerical approximation at 1% accuracy to calculate joint probabilities of the combined discrete classes (Table 2) for total mortality used in the model.

**Table 2.**
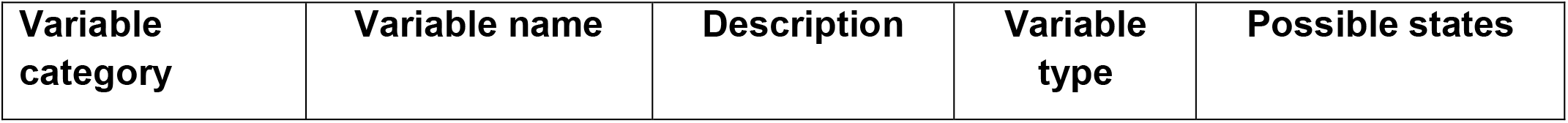
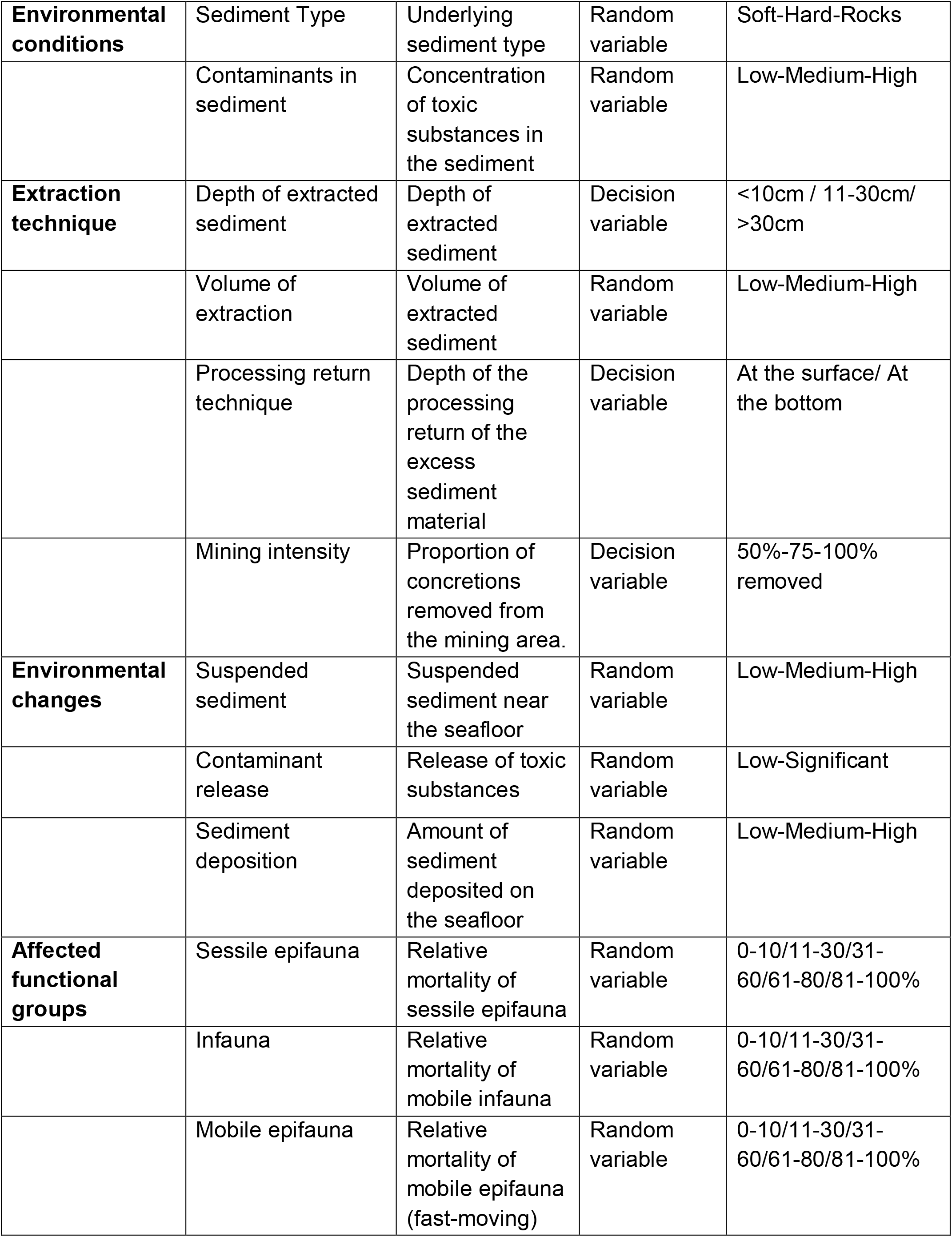
Variables in the Bayesian Network model for ecological risks of seabed mining.

The resulting CPTs were incorporated in the BN model created in R software (R 2020). Using the Bayes rule, BNs enable evaluating different scenarios and to compute posterior probabilities given new knowledge. In this context, a BN allows modification of the operational parameters to evaluate the impacts of different mining operations and the associated changes in the functional groups. The joint probability distribution in the BN may then be used to make queries on the impact of multiple pressures on specific ecosystem components to assess the risks and to evaluate which variables should be monitored to obtain a reasonable overview of the impacts. Here we queried the network on two alternative mining scenarios, which we define as a combination of specific states of the decision variables that describe the overall mining process and are assumed to be controlled by the party responsible for the mining operation (Table 2). The random variables in the model are further affected by these decision nodes (Figure 4, Table 2).

The modelling was done using R 3.6.3, with package *bnlearn* (Scutari 2009). Full details of the model with the R scripts and the conditional probability tables are available at: https://github.com/lkaikkonen/Causal_SBM.

## 4. RESULTS

### 4.1 Causal maps

The expert interviews resulted in 11 individual causal maps. In some cases, the experts took the lead in drawing the variables and connections between them, whereas in most interviews the modeler had the main responsibility of drafting the map based on the discussion.

The number of variables in the individual maps varied between 8 and 24. In general, there were no contradictory views, and the differences between the maps were attributed to the number of variables and level of detail in different processes regarding the impacts of mining. We were not successful in eliciting all link strengths, and only the strongest connections were explicitly given by all experts. The individual causal maps are included in the Supporting Information (S1).

After concept harmonization, the final causal map has 53 variables. Multiple iterations of expert comments on the causal network structure resulted in a combined causal network with 96 connections (Figure 3). The rationale for the connections between variables and further details on them are summarized in Tables S2–S4 in the Supporting Information.

**Figure 3.**
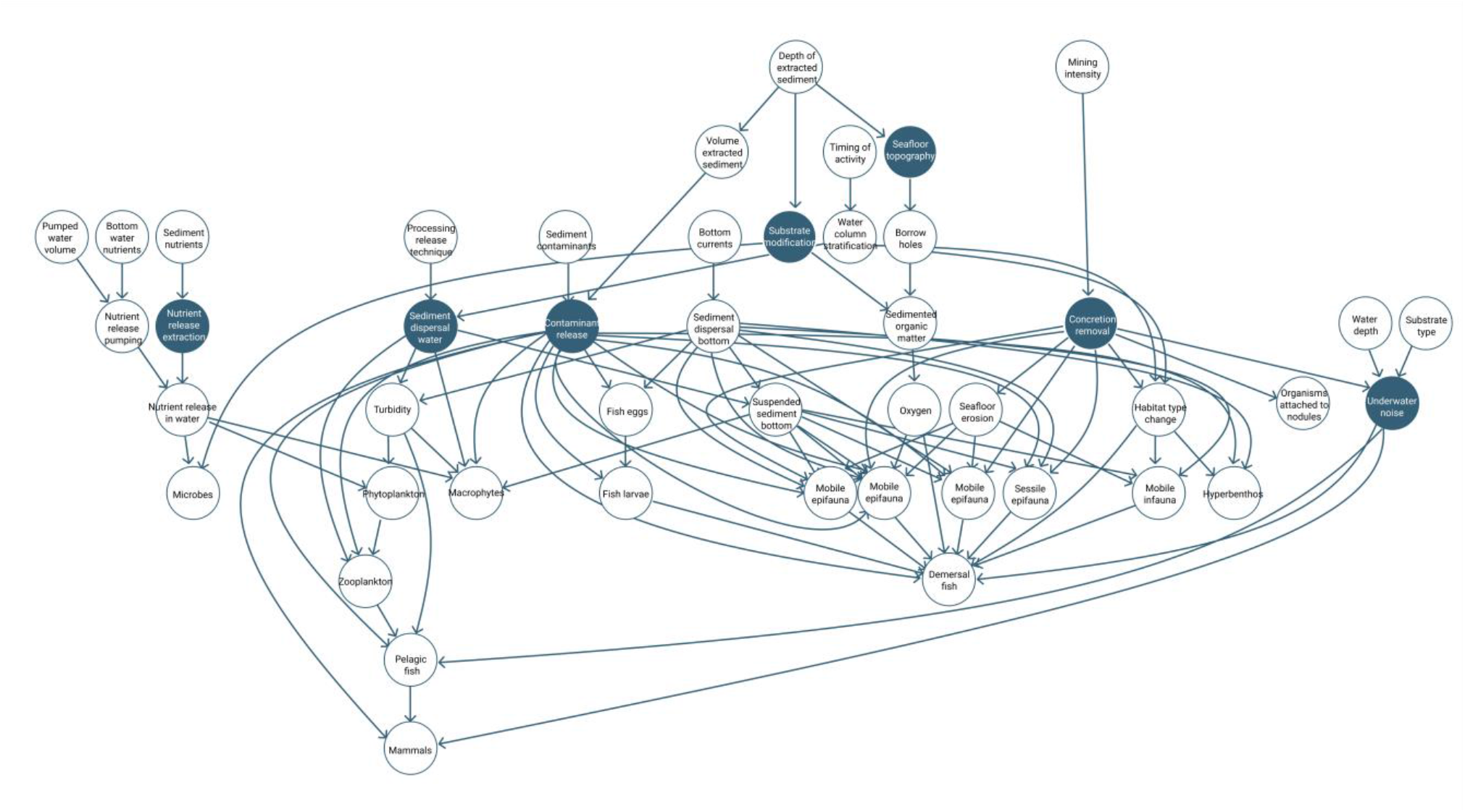
Combined causal map of the environmental and ecological effects of seabed nodule extraction on Baltic Sea ecosystem. The colored ovals denote pressures that were the starting point for each interview and the subsequent causal mapping. For full details of the variables and causal connections, see Tables S2-S4 in the Supporting Information.

### 4.2 Impacts of mining on marine ecosystems: Combined causal network

The first set of interviews with geologists revealed several factors affecting the magnitude of physicochemical changes in the environment, related to both the execution of the mining operation and the prevailing environmental conditions (Table 2). The factors regarding the mining technique included water depth at the extraction site, depth of extracted sediment, and processing return technique. Both the geologists and ecologists included several environmental factors in their causal maps, including variables describing the sediment characteristics and composition, water column chemistry, and hydrological parameters (Figure 3).

The impacts on the biological ecosystem components were more complex and spanned into the spatial and temporal dimensions than the physicochemical changes in the environment. Experts successfully adopted a parsimonious attitude to defining the functional groups and expressed how these groups would be affected by the different pressures. The most detail in terms of functional traits was given to benthic fauna which are most directly affected by substrate extraction. Experts included a wide range of organisms in the assessment that were unlikely directly affected in the extraction area, including early life-stages of fishes, macrophytes, and mammals. Factors affecting the recovery potential of organisms and ecosystem functions after disturbance were mentioned in all interviews.

Direct extraction of seabed substrate and the resulting habitat loss was deemed to have the most significant impact on benthic fauna. Many experts equally considered the impacts of elevated suspended sediment concentrations on filter feeding organisms severe. In the interviews, the functional groups were deemed different in terms of acute impacts of disturbance. For example, while highly mobile organisms like fish are assumed to escape from the extraction area, significant changes in the environment either through modification of bottom substrate or benthic fauna as food are expected to potentially affect the distribution of demersal fish species. Similarly, release of contaminants from the sediment was estimated to significantly affect all organisms, yet it was noted that many toxic effects might only be expressed in the reproductive success of organisms. Nearly all experts noted the negative impacts of underwater noise on mammals and fishes.

### 4.3 Quantitative case study: Acute impacts on benthic fauna

The full causal model is highly complex (Fig. 3), and parameter estimation would be a demanding task. Therefore, for illustration we selected 18 variables for the quantitative analysis to describe the acute impacts on benthic fauna (Figure 4, Table 2). We queried the network on two different mining scenarios. The resulting probability distributions are presented in figure 5.

**Figure 4.**
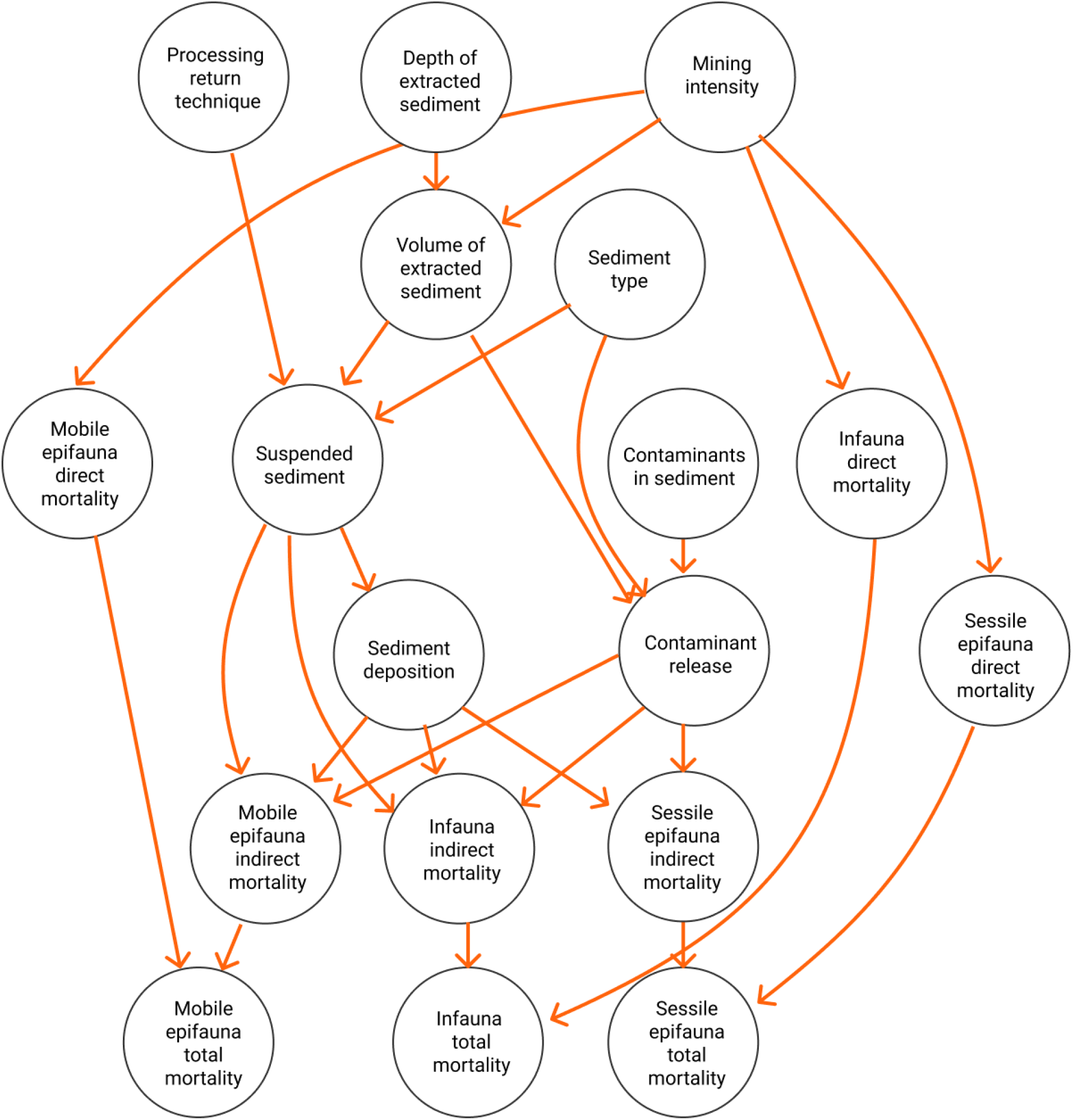
Bayesian network structure for immediate impacts on selected groups of benthic fauna. Mining scenario may be controlled by *processing return technique, depth of extracted sediment*, and *mining intensity*.

In the case of mining 75% of a discrete mining block, the most probable outcome in terms of total mortality for both sessile epifauna and infauna is estimated to be 81–100% mortality (Fig. 5,A). The probability of the highest mortality for sessile epifauna is slightly higher than for infauna (60.1% compared to 57.7%, respectively). For mobile epifauna, 60–80% mortality is the most likely outcome with a 52.2% probability.

**Figure 5.**
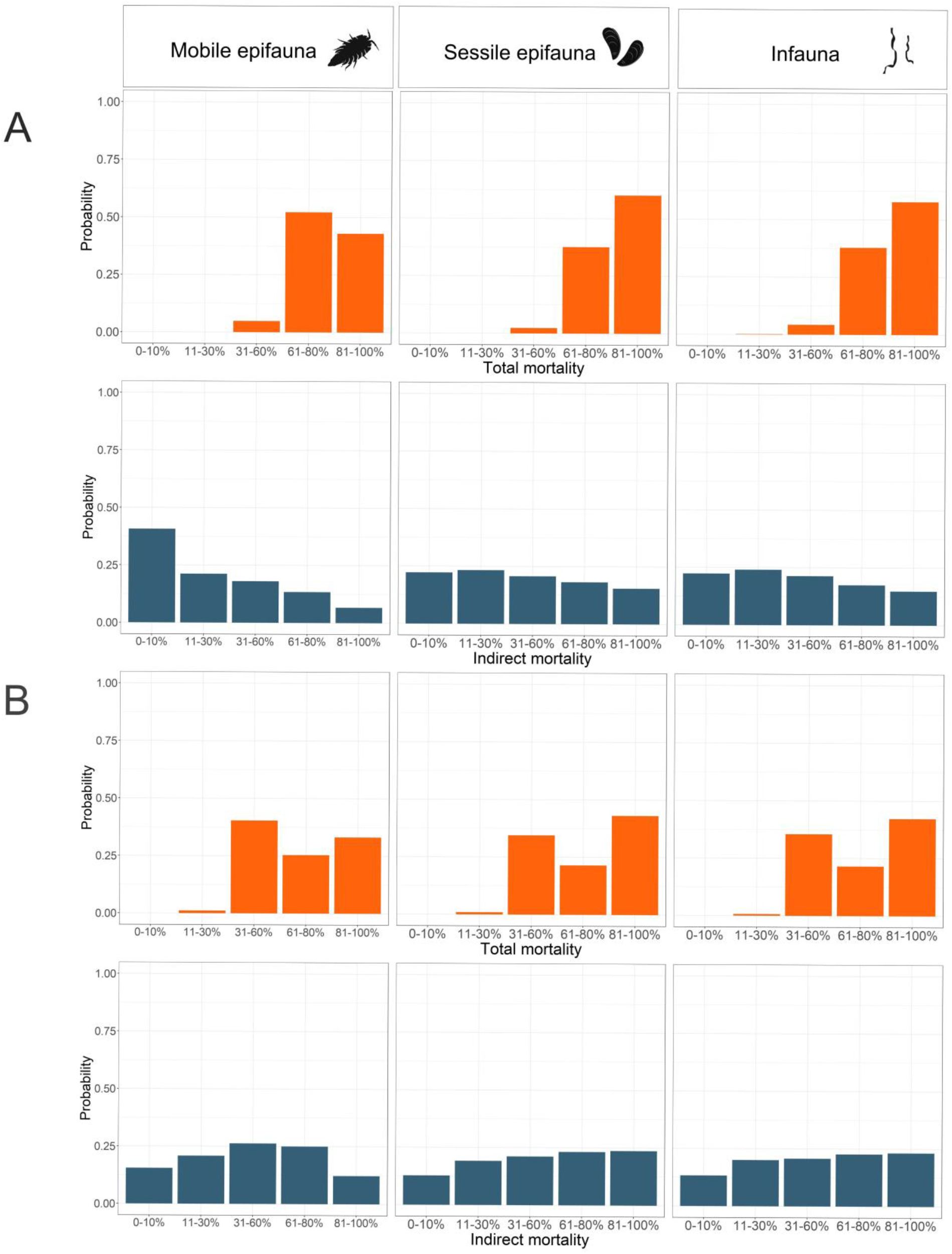
Joint probability distribution of the total and indirect mortality of mobile epifauna, sessile epifauna, and infauna under two alternative mining scenarios: A) Mining 75% of a discrete mining block with 11-30cm sediment extracted, and B) mining 50% of a discrete mining block with 11-30cm sediment extracted with release of harmful substances from the sediment.

The likeliest outcome of the mining scenario described above in terms of indirect mortality resulted in indirect mortality of 11–30% of both infauna (24.1% probability) and sessile epifauna (23.3% probability) and 0-10% mortality of mobile epifauna with 40.7% probability (Fig. 5, A). The probability of the highest mortality (81–100%) is 14.8% for infauna, 15.5% for sessile epifauna and 6.6% for mobile epifauna. Overall, the probability of both indirect and direct mortality on sessile epifauna and infauna are deemed equally widely distributed.

The BN model allows estimating the probability of any variable of interest in the model (here relative mortality) given certain evidence (e.g. regarding the mining operation or environmental conditions). To give an example, when mining occurs on only 50% of a discrete block, but release of harmful substances is known to occur, the probabilities for the indirect mortality of benthic fauna are higher for all groups (Fig. 5,B). These changes illustrate the relative importance of certain pressures on the overall mortality.

Changes in the extent of direct extraction of seabed substrate and FeMn concretions had the largest impact on the direct mortality of the benthic fauna. In terms of indirect effects, the release of ecologically significant levels of toxic substances from the sediment had the highest impact on the mortality of benthic fauna. In a similar way, the model may be used to evaluate the cumulative effects of multiple stressors for each assessed ecosystem component by first ranking the relative effects of each stressor on the mortality of the community and then evaluating the probability distribution for each combination of stressor levels.

## 5. DISCUSSION

This study presents the first systematic evaluation of the ecological risks associated with seabed mining. By interviewing a multidisciplinary group of experts, we outline a basis for an ecological risk assessment model. We further demonstrate how qualitative information may be used to move towards a quantitative assessment by using a causal probabilistic approach to estimate the impacts of seabed disturbance and direct sediment extraction on benthic fauna in the Baltic Sea. These results show that the knowledge related to the impacts of seabed mining is still low, calling for further research on the risks of mining if the operation permits are to be based on a valid scientific understanding.

Involving multiple experts in consecutive interviews provided a comprehensive view of the pressures arising from mining, factors affecting the magnitude of the physicochemical changes, and the affected ecosystem components. Particularly the interviews with geologists enabled the inclusion of operational variables related to mining activity and environmental conditions that were deemed to govern the magnitude of pressures. Most detail in terms of affected biological components was given to benthic faunal groups from all ecologists. While we had expected experts to prioritize their own fields’ species in more detail, this was not always the case, and the experts’ previous participation in similar mapping exercises seemed to be the factor governing the number of connections and variables.

Although many of the impact pathways described in the obtained causal maps have been cited in previous studies (Koschinsky et al. 2018; Christiansen et al. 2020), our mapping exercise enabled a more detailed inclusion of pelagic ecosystem components which have been neglected in many previous studies on seabed impacts (Newell et al. 2004; Boyd et al. 2005; Krause et al. 2010; Christiansen et al. 2020). A qualitative causal representation of the impacts alone can thus help better understand how risks emerge and can potentially be mitigated (Chen and Pollino 2012; Carriger et al. 2018). Drafting the causal maps from the beginning further ensures that all relevant connections are included, and biases in thinking will be revealed easier (Tversky and Kahneman 1979; Renn 2008).

Depending on the extraction intensity and the functional group, acute mortality of benthic fauna was estimated to be most likely at rates of 60–100% in the directly affected area and 0–10% to 10–30% in the indirectly affected area. The probabilities of very high indirect mortality (81–100%) were over 10% in both of the evaluated scenarios for sessile epifauna and infauna. Accounting for the indirect mortality separately allows further refining the assessment to account for the impacts of indirect effects, as these are deemed significant in terms of the spatial footprint due to dispersal of suspended sediment (Boyd and Rees 2003; Desprez et al. 2009).

Overall, the probability distributions on the relative mortality of benthic fauna from expert assessment are rather broad, showing low levels of certainty on the impacts. One reason for this is likely the lack of scientific knowledge, particularly regarding the cumulative effects from multiple pressures, which make validating such assessments challenging. Although the different functional groups of benthic fauna were deemed to experience differential responses particularly due to indirect impacts from sediment deposition and suspended sediment, the probability distributions describing these effects are very similar between infauna and sessile epifauna. While these results may be a consequence of the high uncertainties related to the impacts, further knowledge engineering approaches to facilitate elicitation (Martin et al. 2012; Laitila and Virtanen 2016) could offer insights into the effects of multiple pressures. Future development of the model should thus address improving the quantitative estimates of the risks in terms of both methodology and the used evidence

### Expert knowledge in ecological risk assessments

The interviews and the subsequent causal mapping highlighted the challenges in conceptualizing spatiotemporal complexity related to anthropogenic impacts (Gladstone-Gallagher et al. 2019). Although we had specifically requested experts to focus on a discrete spatially defined area and immediate impacts, factors affecting recovery and spatial extent of impacts arose in all interviews. These differences in temporal scale are a result of changes in the environment varying in their scope and persistence (see Table S5 for spatial and temporal extent of the pressures), resulting in immediate impacts, chronic and long-term impacts, and factors affecting the recovery potential of organisms. To operationalize a multidimensional view of risks and to move towards a quantitative assessment, it is necessary to consider which pressures operate at which time scales and spatial dimensions.

Given these challenges, attempting direct modelling of such dynamic systems may not be appropriate, as it can result in excessive simplification and loss of information. Giving the experts free hands was beneficial for capturing also the non-immediate impacts and in retrospective, our interviews could have been developed in a more flexible manner. We argue, however, that providing starting points for the assessment by setting the spatial and temporal limits helped the experts to get started without being tangled in the multidimensionality. The results show that it is essential to consider effects from multiple perspectives and account for the multidimensional disturbance space. An operational assessment should thus include multiple time steps or account for continuous effects and changes in the prevailing conditions.

### How can predictive risk assessment inform marine resource governance?

The paucity of evidence on the impacts of seabed mining calls for more comprehensive views of the risks and knowledge gaps to support decision-making. Given the modular structure of BNs, the model presented here may be adapted for more complex ERA through separate layers and sub-models. While this model provides only a limited view of the relationships within food webs, functional ecology and biogeochemical connections, it is a starting point for more detailed ecological risk assessments. Another advantage of probabilistic approaches is that the conditional probabilities may be drawn from multiple sources and can include both qualitative and quantitative data. This allows iterative updating of the model as new information becomes available. BNs can further be developed into dynamic networks that can also account for temporal changes to measure resilience and recovery of ecosystems (Wu et al. 2018).

To support decision-making on potential future use of seabed resources and further evaluation of trade-offs from mining, model simulations under alternative mining scenarios should be compared to existing policy targets regarding acceptable changes in ecosystems. Using a quantitative approach offers a more robust and transparent means of estimating the impacts of emerging activities when defining acceptable thresholds to the impacts (Levin et al. 2016). With recent calls for more empirical approaches to the broad scale seabed mining initiatives (Drazen et al. 2020 Jul 8), new data on the impacts of mining may be incorporated in the risk model to learn the probability distributions between the nodes from data, and further be completed with expert assessment. Estimating the impacts and accounting for the knowledge gaps with a probabilistic approach can aid to either support a moratorium and not to go ahead with exploitation in line with a precautionary approach (Barbier et al. 2014), or to provide information for more comprehensive risk management plans for potential future mining activities, including the need for mitigation measures. In a case where uncertainties are considered too high, permits could be made to be conditional on improved knowledge by allowing only one mining operation to proceed until impacts have been documented in more detail(Smith et al. 2020), urging the industry to carry out further studies.

Causal networks may be enhanced into more comprehensive frameworks for integrated environmental assessments to promote deeper engagement of multiple values and stakeholders in policy-making (Mourhir et al. 2016). Using a systematic framework with causal networks helps paint a more complete picture of the system and the associated environmental impacts, enabling better inclusion of uncertainty in the environmental management plans of seabed resource use and improving transparency of the estimates. Engaging with multiple experts and sources of knowledge not only strengthens the knowledge base for assessing the risks, but also allows revealing possibly contradictory views between experts and stakeholders (Freudenburg et al. 1999).

The expanding industrial use of the ocean space and resources calls for more detailed assessments on the risks associated with them. Recent incentives for more sustainable marine governance (Lubchenco et al. 2016; Golden et al. 2017; Bennett et al. 2019) further urge applying an ecosystem approach to resource management, including impact and risk assessments of activities on both the marine ecosystem and human society.

Based on the results of this study, we posit that while empirical observations are key in unravelling the impacts of novel activities, full consideration of the different scales of risks requires a systematic approach to bring together findings from empirical studies, modelling, and expert assessments. An improved view of the risks as an underlying concept in research on the impacts of seabed mining will aid developing integrative ecosystem based management of emerging maritime industries (Hodgson et al. 2019).

## ACKNOWLEDGEMENTS

This research was funded by the Strategic Research Council at the Academy of Finland, under projects SmartSea (grant number 292?985) and WISE (grant number 312627). It was also funded by the Academy of Finland grant number 311944 to the strategic research profiling area The Sea at åbo Akademi University (AT), well as through the BONUS BLUEWEBS project which has received funding from BONUS (Art 185), funded jointly by the EU and the Academy of Finland (LU). IH was funded by the Helsinki Institute of Sustainability Science (HELSUS), University of Helsinki. The authors sincerely thank all the experts involved in this study for kindly providing their time and expertise.

## AUTHOR CONTRIBUTIONS

[CRediT taxonomy:](https://casrai.org/credit/)

Conceptualization: LK; SK, RV; Methodology: LK, LU, IH; Formal analysis and investigation: LK; Writing - original draft preparation: LK, Writing - review and editing: AT, HN, LU, IH, KK, RV, SK; Funding acquisition: SK; Supervision: LU, IH, KK, SK, RV

## SUPPORTING INFORMATION

Supporting information (SI S1-S5) are available as an attachment to this manuscript, as well as at https://github.com/lkaikkonen/Causal_SBM.

