## SupplementaryMaterial for "Causal approach to environmental risks of seabed mining" (from another version of this preprint: version 1, 2021-02-21)

### SUPPLEMENTARY MATERIAL

*This file contains the supporting information for Kaikkonen, L. et. al.: Causal Approach to Environmental Risks of Seabed Mining*

**S1** Complete causal maps from interviews. The colored ovals depict the pressures which were presented at the beginning of the interviews and served as a starting point for the causal mapping exercise.

#### Geologists

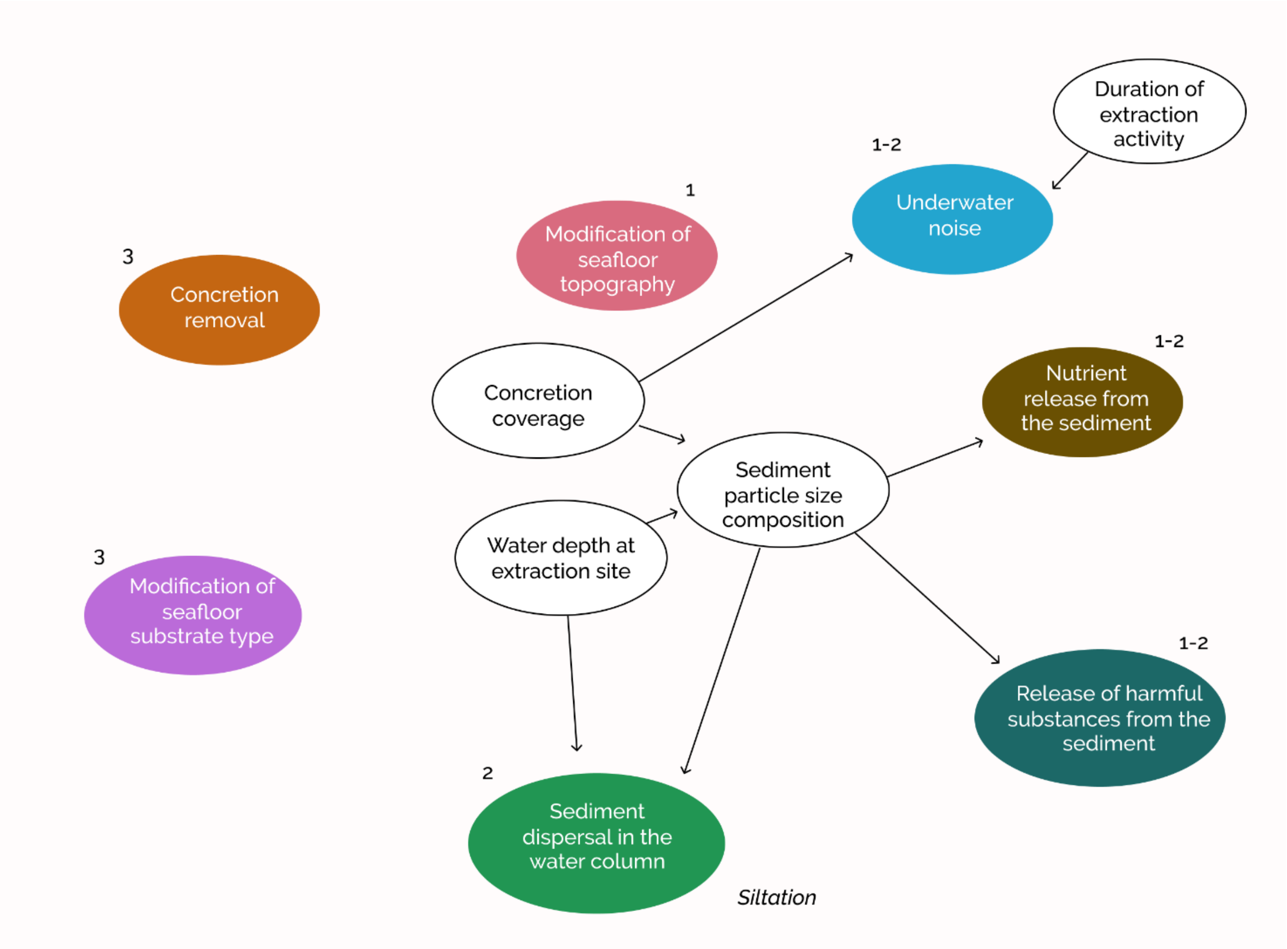

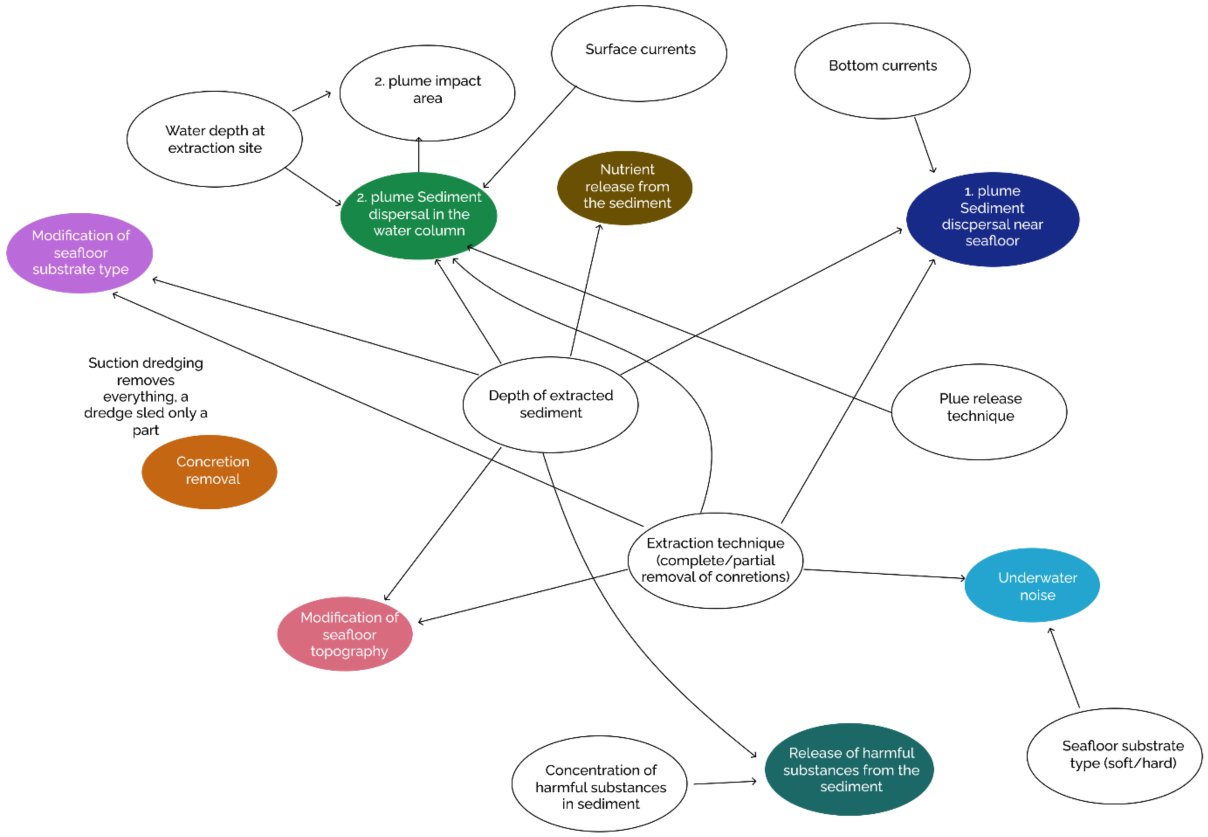
Surface currents
Bottom currents
2. plume impact area
Water depth at extraction site
2. plume Sediment dispersal in the water column
Nutrient release from the sediment
1. plume Sediment discpersal near seafloor
Modification of seafloor substrate type
Suction dredging removes everything, a dredge sled only a part
Concretion removal
Depth of extracted sediment
Plue release technique
Extraction technique (complete/partial removal of conretions)
Underwater noise
Modification of seafloor topography
Seafloor substrate type (soft/hard)
Concentration of harmful substances in sediment
Release of harmful substances from the sediment

9

10

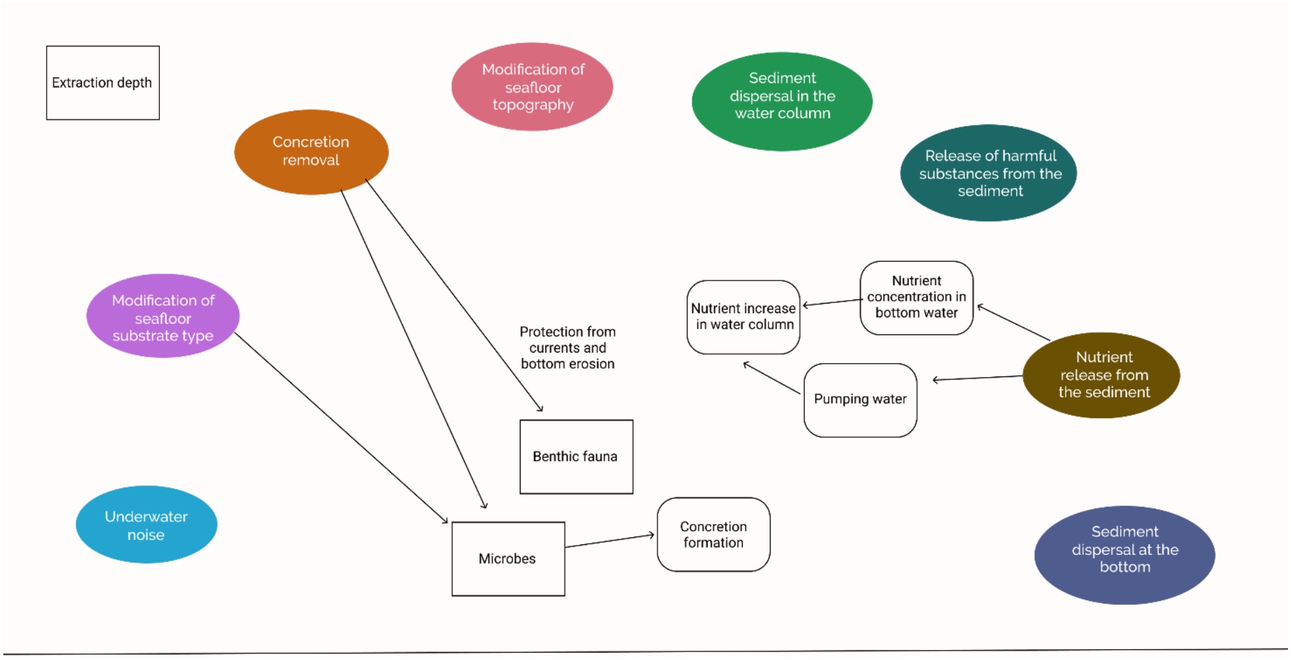
Extraction depth
Modification of seafloor topography
Sediment dispersal in the water column
Concretion removal
Release of harmful substances from the sediment
Modification of seafloor substrate type
Nutrient concentration in bottom water
Nutrient increase in water column
Protection from currents and bottom erosion
Nutrient release from the sediment
Pumping water
Benthic fauna
Underwater noise
Microbes
Concretion formation
Sediment dispersal at the bottom

11

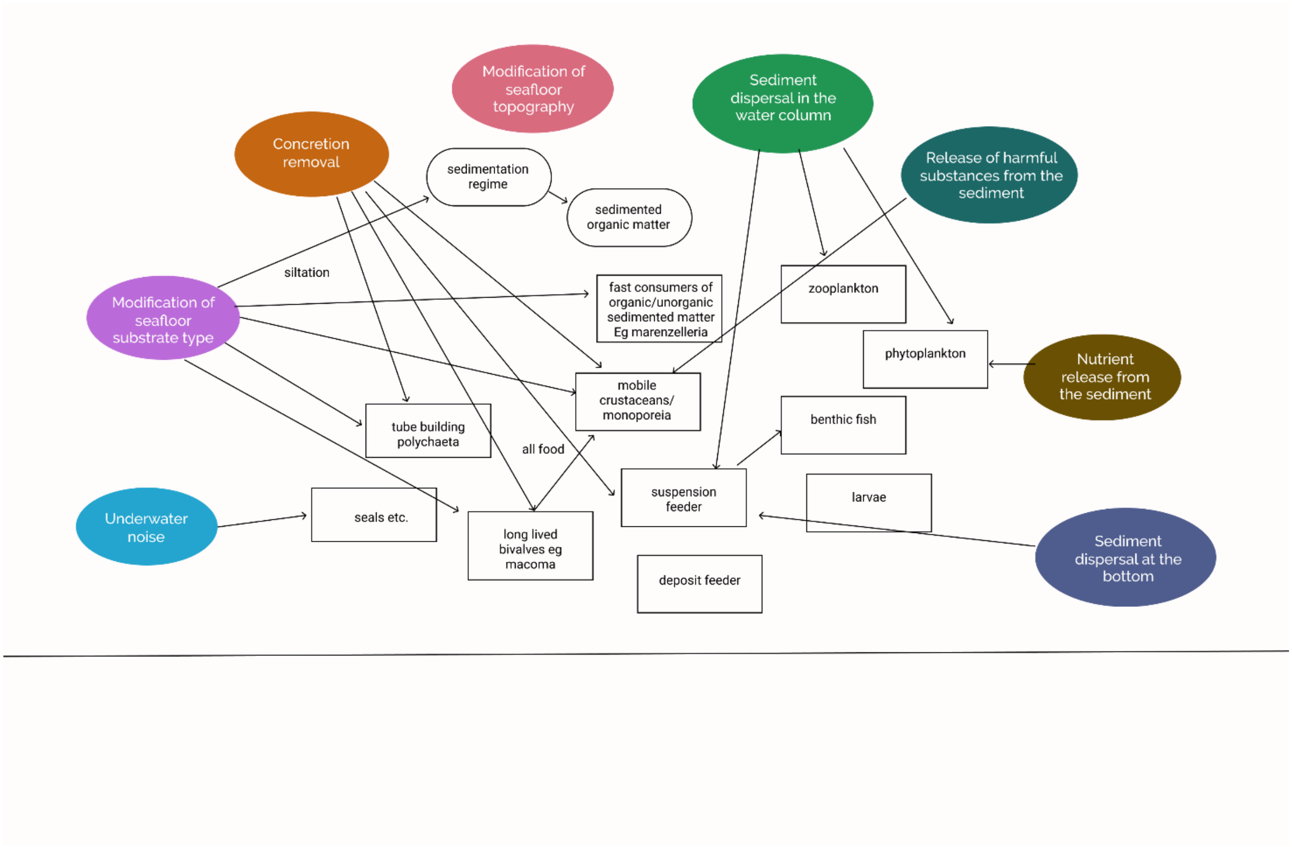
Modification of seafloor topography
Sediment dispersal in the water column
Concretion removal
Release of harmful substances from the sediment
sedimentation regime
sedimented organic matter
siltation
Modification of seafloor substrate type
fast consumers of organic/unorganic sedimented matter Eg marenzelleria
zooplankton
phytoplankton
Nutrient release from the sediment
mobile crustaceans/ monoporeia
tube building polychaeta
benthic fish
all food
suspension feeder
larvae
Underwater noise
seals etc.
long lived bivalves eg macoma
Sediment dispersal at the bottom
deposit feeder

12

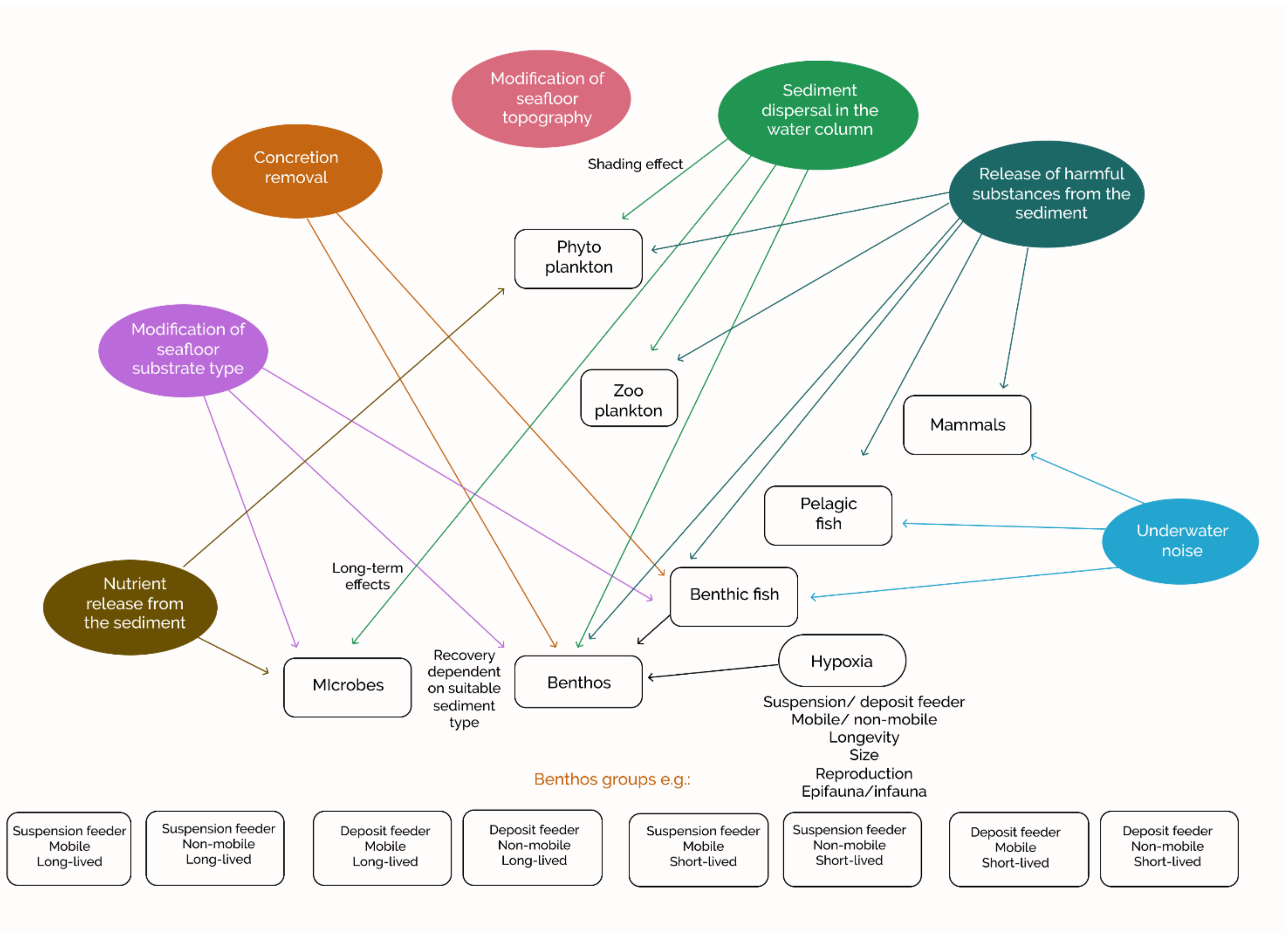
Modification of seafloor topography
Sediment dispersal in the water column
Concretion removal
Shading effect
Release of harmful substances from the sediment
Phyto plankton
Modification of seafloor substrate type
Zoo plankton
Mammals
Pelagic fish
Underwater noise
Long-term effects
Benthic fish
Nutrient release from the sediment
Recovery dependent on suitable sediment type
Hypoxia
MIcrobes
Benthos
Suspension/ deposit feeder
Mobile/ non-mobile
Longevity
Size
Reproduction
Epifauna/infauna
Benthos groups e.g.:
Suspension feeder Mobile Long-lived
Suspension feeder Non-mobile Long-lived
Deposit feeder Mobile Long-lived
Deposit feeder Non-mobile Long-lived
Suspension feeder Mobile Short-lived
Suspension feeder Non-mobile Short-lived
Deposit feeder Mobile Short-lived
Deposit feeder Non-mobile Short-lived

13

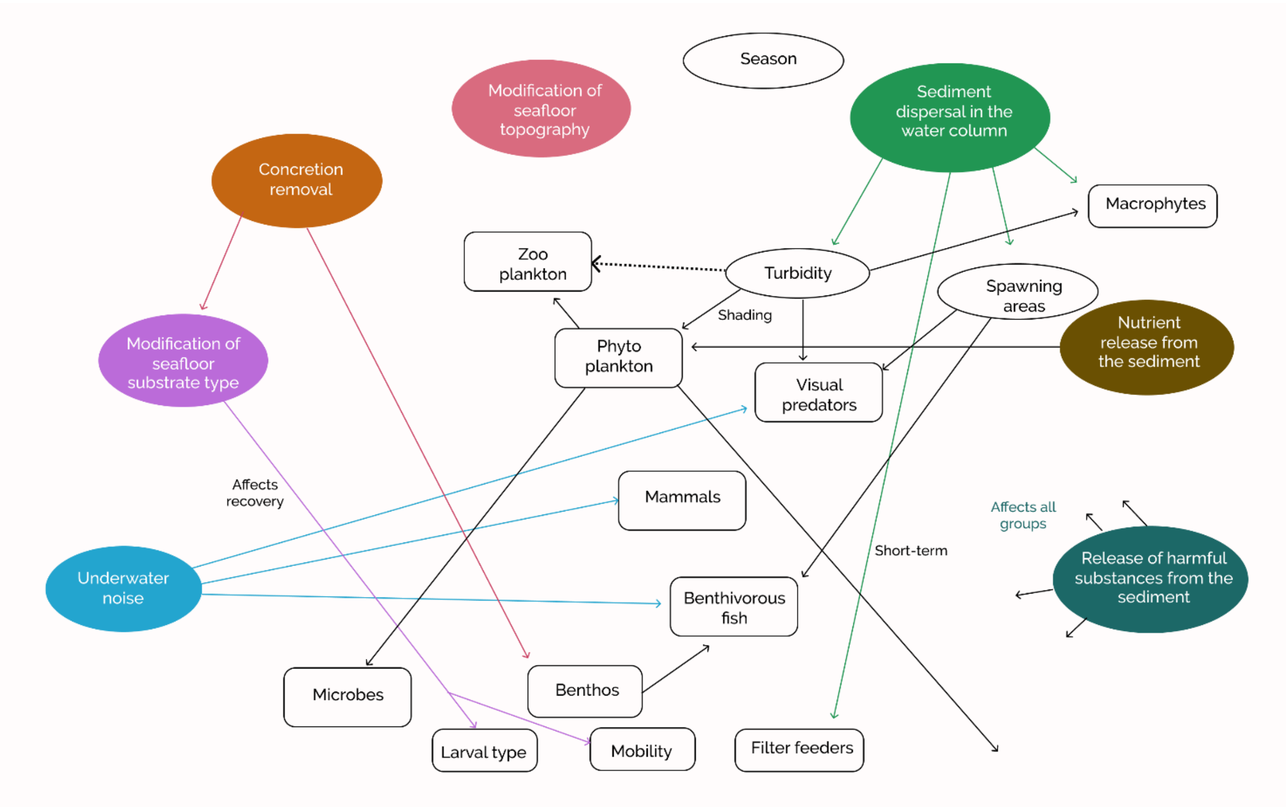
Season
Modification of seafloor topography
Sediment dispersal in the water column
Concretion removal
Macrophytes
Zoo plankton
Turbidity
Spawning areas
Shading
Phyto plankton
Nutrient release from the sediment
Modification of seafloor substrate type
Visual predators
Affects recovery
Mammals
Affects all groups
Short-term
Release of harmful substances from the sediment
Underwater noise
Benthivorous fish
Microbes
Benthos
Larval type
Mobility
Filter feeders

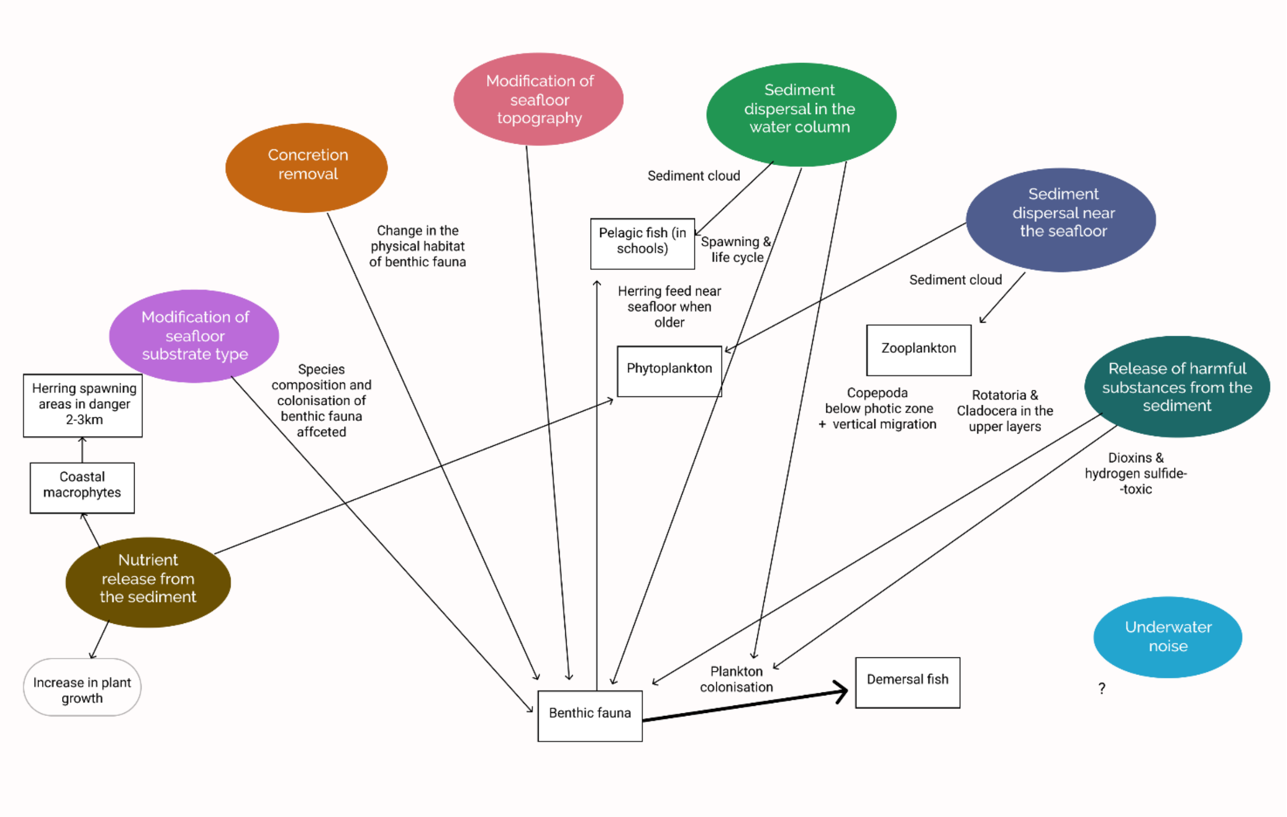
Modification of seafloor topography
Sediment dispersal in the water column
Concretion removal
Sediment cloud
Pelagic fish (in schools)
Spawning & life cycle
Change in the physical habitat of benthic fauna
Sediment dispersal near the seafloor
Herring feed near seafloor when older
Sediment cloud
Modification of seafloor substrate type
Zooplankton
Phytoplankton
Species composition and colonisation of benthic fauna affceted
Herring spawning areas in danger 2-3km
Copepoda below photic zone + vertical migration
Rotatoria & Cladocera in the upper layers
Release of harmful substances from the sediment
Coastal macrophytes
Dioxins & hydrogen sulfide--toxic
Nutrient release from the sediment
Underwater noise
Increase in plant growth
Plankton colonisation
Demersal fish
Benthic fauna
?

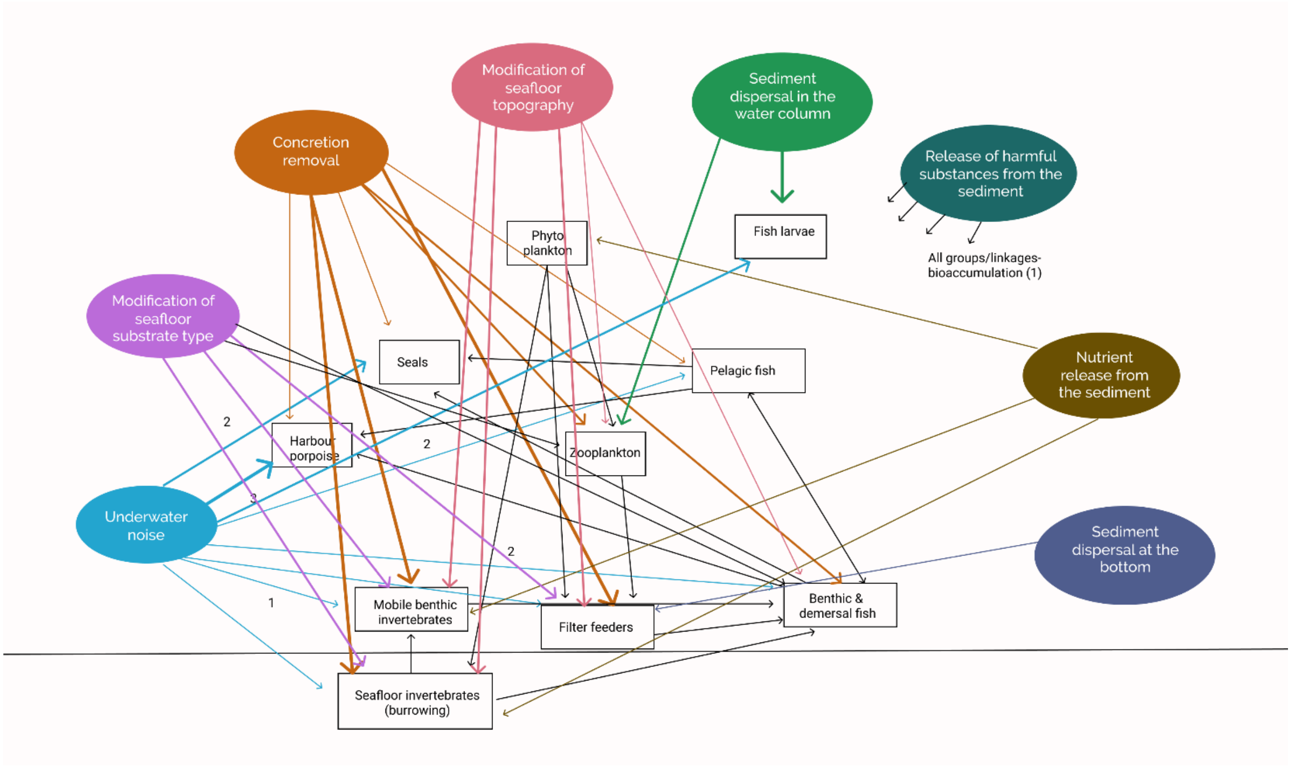
Modification of seafloor topography
Sediment dispersal in the water column
Concretion removal
Release of harmful substances from the sediment
Fish larvae
Phyto plankton
All groups/linkages- bioaccumulation (1)
Modification of seafloor substrate type
Seals
Pelagic fish
Nutrient release from the sediment
2
Harbour porpoise
2
Zooplankton
Underwater noise
3
2
Sediment dispersal at the bottom
Mobile benthic invertebrates
Benthic & demersal fish
1
Filter feeders
Seafloor invertebrates (burrowing)

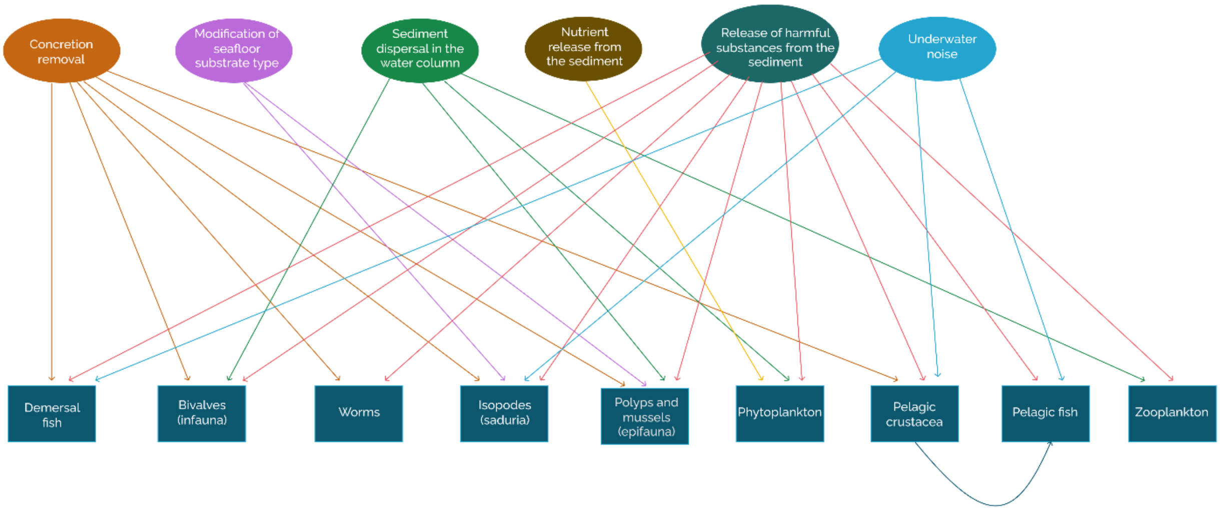
Concretion removal
Modification of seafloor substrate type
Sediment dispersal in the water column
Nutrient release from the sediment
Release of harmful substances from the sediment
Underwater noise
Demersal fish
Bivalves (infauna)
Worms
Isopodes (saduria)
Polyps and mussels (epifauna)
Phytoplankton
Pelagic crustacea
Pelagic fish
Zooplankton

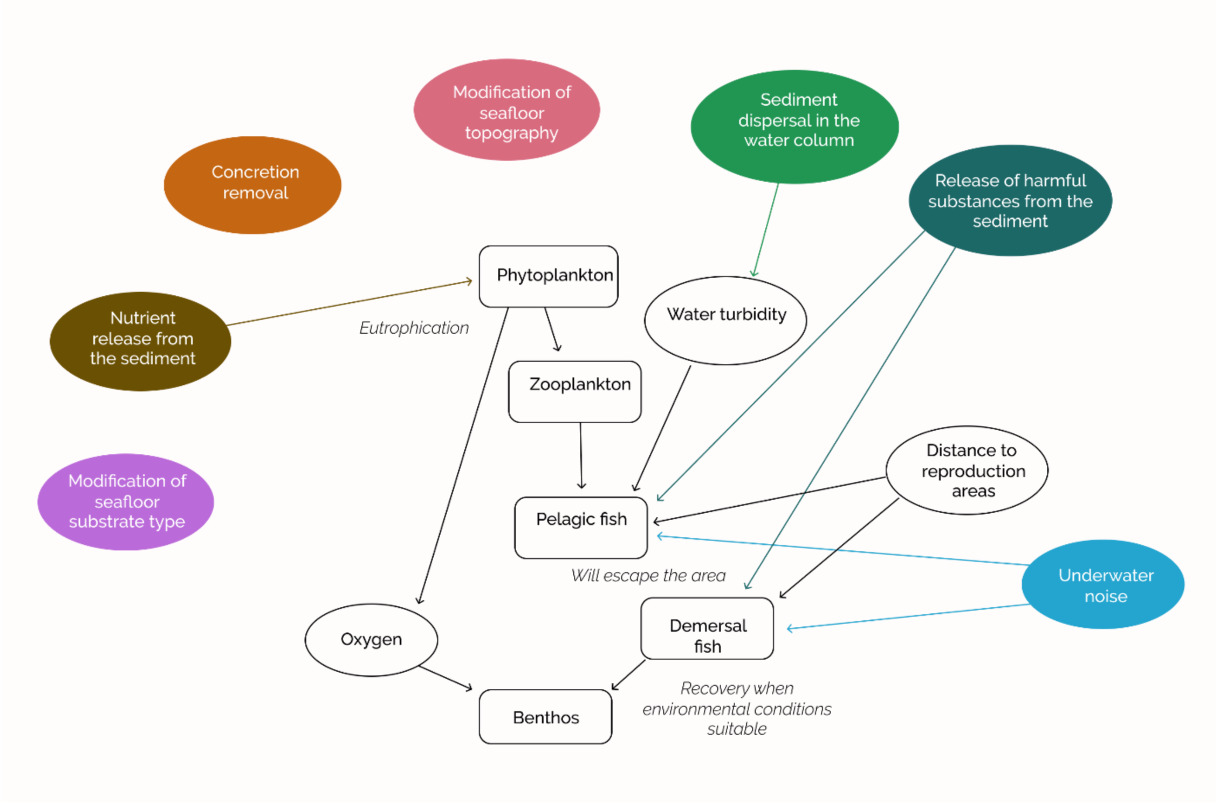
Modification of seafloor topography
Sediment dispersal in the water column
Concretion removal
Release of harmful substances from the sediment
Phytoplankton
Water turbidity
Nutrient release from the sediment
Eutrophication
Zooplankton
Modification of seafloor substrate type
Distance to reproduction areas
Pelagic fish
Underwater noise
Will escape the area
Demersal fish
Oxygen
Benthos
Recovery when environmental conditions suitable

22

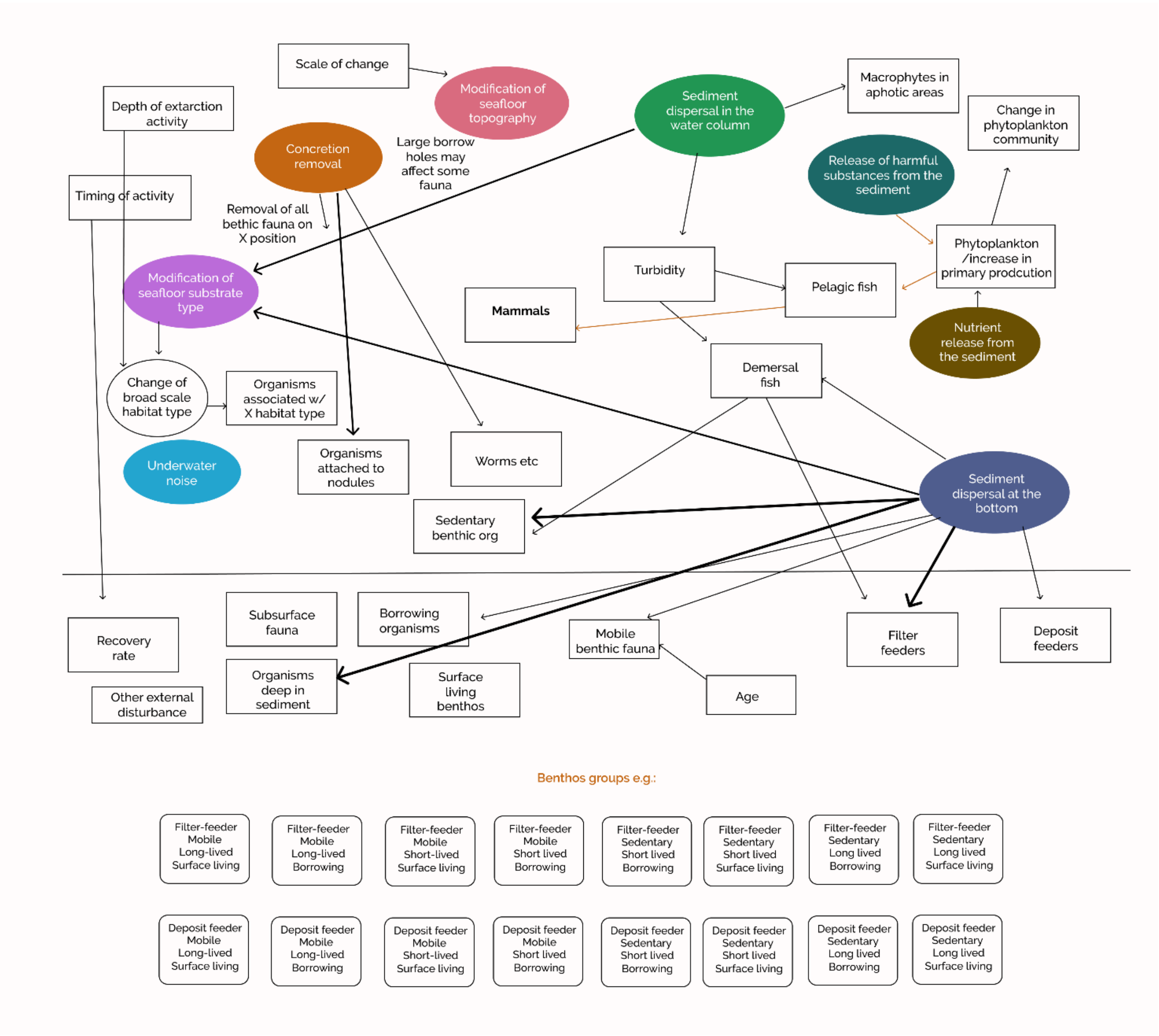


**Table S2.** Causal interactions between ecosystem components & pressures affecting them.

| Functional groups of organisms | Affecting pressures or other variables |
|---|---|
| **Mammals (porpoises & seals)** | **1.** Underwater noise induces stress.<br>**2.** Harmful substances may reduce fitness in mammals.<br>**3.** Quantity and quality of fish as food source affect the number of mammals in the area. |
| **Demersal fish** | **1.** Underwater noise induces stress & causes fish to migrate away. Migration requires energy, which reduces the fitness of individuals.<br>**2.** Sediment dispersal in the water column & on the seafloor may cause |

| | |
|---|---|
| | fish to avoid extraction area during activity.<br>**3.** Harmful substances reduce fitness in fish, which can be reflected in the reproductive success of individuals.<br>**4.** Change in habitat type may cause fish to leave area.<br>**5.** Anoxia causes fish to avoid extraction area.<br>**6.** Quantity of benthic invertebrates as food affects the number of demersal fish in the area. |
| **Pelagic fish** | **1.** Underwater noise induces stress & can cause fish to avoid extraction area.<br>**2.** Zooplankton biomass as food affects number of pelagic fish in the area.<br>**3.** Number of fish larvae affects abundance of adult fish.<br>**4.** Turbidity affects predation success, reduces fitness & can cause fish to avoid extraction area. |
| **Fish eggs** | **1.** Sedimentation may blanket eggs.<br>**2.** Harmful substances may destroy eggs. |
| **Fish larvae** | **1.** Bottom sedimentation may smother larvae.<br>**2.** Underwater noise may reduce fitness.<br>**3.** Harmful substances may kill larvae.<br>**4.** Quantity of fish eggs affects larval abundance. |
| **Phytoplankton** | **1.** Sediment in the water column may shade phytoplankton.<br>**2.** Toxin release may modify community composition.<br>**3.** Nutrients increase primary production. |
| **Zooplankton** | **1.** Sediment in water column may clog feeding organs & impact individual fitness.<br>**2.** Quantity and quality of phytoplankton as food source affects zooplankton biomass. |
| **Pelagic crustacean** | **1.** Underwater noise induces stress and reduces fitness in organisms.<br>**2.** Toxic substances may reduce fitness.<br>**3.** Quantity and quality of mobile infauna as a food source affect biomass.<br>**4.** Habitat modification may affect recolonization capacity. |
| **Macrophytes** | **1.** Higher nutrient concentrations increase macrophytes growth. |

| | |
|---|---|
| | **2.** Turbidity reduces light availability in the water column and changes the quality of light.<br>**3.** Toxic substances may affect macrophyte growth.<br>**4.** Sedimentation may smother macrophytes. |
| **Benthic microbes** | **1.** Removal of specific taxa through concretion removal.<br>**2.** Nutrient release from sediment affects growth. |
| **Organisms attached to nodules** | **1.** Concretion extraction removes organisms.<br>**2.** Sediment plume may clog feeding organs & blanket organisms |
| **Sessile epifauna filter/suspension feeder** | **1.** Noise may induce stress & reduce fitness.<br>**2.** Concretion extraction removes organisms.<br>**3.** Sediment plume may clog and blanket organisms.<br>**4.** Low oxygen concentrations reduce fitness.<br>**5.** Toxic substances reduce fitness. |
| **Mobile infauna filter/suspension / deposit feeder** | **1.** Noise may induce stress.<br>**2.** Concretion extraction removes organisms.<br>**3.** Habitat change affects recovery<br>**4.** Sediment plume may clog feeding organs.<br>**5.** Low oxygen concentrations reduce fitness.<br>**6.** Toxic substances reduce fitness. |
| **Mobile epifauna filter / suspension feeder** | **1.** Noise may induce stress<br>**2.** Concretion extraction removes organisms.<br>**3.** Habitat change affects recovery<br>**4.** Sediment plume may blanket<br>**5.** Low oxygen concentrations reduce fitness. |
| **Mobile epifauna deposit-feeder (slow-moving)** | **1.** Noise may induce stress.<br>**2.** Concretion extraction removes organisms.<br>**3.** Habitat change affects recovery.<br>**4.** Sediment plume may blanket organisms.<br>**5.** Low oxygen concentrations reduce fitness.<br>**6.** Toxic substances reduce fitness. |
| **Mobile epifauna predator (fast-moving)** | **1.** Noise may induce stress. |

| | |
|---|---|
| | **2.** Habitat change affects recovery<br>**3.** Sediment plume may blanket organisms.<br>**4.** Low oxygen concentrations reduce fitness.<br>**5.** Toxic substances reduce fitness. |

**Table S3.** Causal connections between the physicochemical pressures and other model variables

| Pressures from mineral extraction | Parameters affecting pressures |
|---|---|
| **Underwater noise** | **1.** Seafloor substrate type (hard/soft) affects noise levels.<br>**2.** Timing of activity (affects stratification in the water column). |
| **Concretion removal** | **-** |
| **Modification of seafloor substrate type** | **1.** Depth of extracted sediment.<br>**2.** Sediment dispersal at the bottom. |
| **Modification of seafloor topography** | **1.** Depth of extracted sediment |
| **Sediment dispersal in the water column** | **1.** Water column stratification<br>**2.** Surface currents |
| **Sediment dispersal at the bottom** | **1.** Depth of extracted sediment affects amount of sediment dispersed<br>**2.** Bottom currents affect sediment dispersal to neighboring areas.<br>**3.** type of extracted sediment |
| **Release of harmful substances from the sediment** | **1.** Concentration of toxic substances in the sediment affects release. |
| **Nutrient release from sediment** | - nutrient concentrations in the sediment |
| **Nutrient release from pumping up bottom water** | **1.** Bottom water nutrient concentrations.<br>**2.** Volume of pumped water. |

**Table S4.** Causal connection between other model variables

| Model variable | Factors affecting variable |
|---|---|
| **Turbidity in the euphotic zone** | **1.** Sediment dispersal in the water column increases turbidity |
| **Turbidity at the bottom** | **1.** Sediment dispersal in the water column<br>**2.** Sediment dispersal at the bottom. |
| **Sediment concentration at the bottom** | **1.** Sediment dispersal at the bottom. |

| | |
|---|---|
| **Change in habitat type** | **1.** Concretion removal<br>**2.** Modification of seafloor substrate type<br>**3.** Sediment dispersal at the bottom. |
| **Borrow holes** | **1.** Depth of extracted sediment affects creation of borrow holes. |
| **Seafloor erosion** | **1.** Concretion removal increases seafloor erosion when the hard substrates and cover is removed. |
| **Nutrient increase in water** | **1.** Nutrient release from sediment<br>**2.** Nutrient release from pumping up bottom water. |

**Table S5.** Spatiotemporal extent of the stressors.

| **Stressor** | **Spatial extent** | **Temporal extent** |
|---|---|---|
| Habitat loss | Local | Long-term |
| Sediment deposition | Local to Regional | Transient to Long-term |
| Sediment substrate type | Local | Long-term |
| Suspended sediment | Local to Regional | Transient |
| Contaminant release | Local to Regional | Long-term |
| Nutrient increase | Regional | Transient |
| Underwater noise | Regional | Transient |
